# Heterogeneous susceptibility to rotavirus infection and gastroenteritis in two birth cohort studies: parameter estimation and epidemiological implications

**DOI:** 10.1101/242172

**Authors:** Joseph A. Lewnard, Benjamin A. Lopman, Umesh D. Parashar, Aisleen Bennett, Naor Bar-Zeev, Nigel A. Cunliffe, Prasanna Samuel, M. Lourdes Guerrero, Guillermo Ruiz-Palacios, Gagandeep Kang, Virginia E. Pitzer

**Author notes:** The findings and conclusions in this report are those of the authors and do not necessarily represent the official position of the Centers for Disease Control and Prevention.

## Abstract

Variation in susceptibility is a known contributor to bias in studies estimating immune protection acquired from vaccination or natural infection. However, difficulty measuring this heterogeneity hinders assessment of its influence on estimates. Cohort studies, randomized trials, and post-licensure studies have reported reduced natural and vaccine-derived protection against rotavirus gastroenteritis in low- and middle-income countries (LMICs). We sought to understand differences in susceptibility among children enrolled in two birth-cohort studies of rotavirus in LMICs, and to explore the implications for estimation of immune protection. We re-analyzed data from studies conducted in Mexico City, Mexico and Vellore, India. Cumulatively, 573 unvaccinated children experienced 1418 rotavirus infections and 371 episodes of rotavirus gastroenteritis (RVGE) over 17,636 child-months. We developed a model characterizing susceptibility to rotavirus infection and RVGE among children, accounting for aspects of the natural history of rotavirus and differences in transmission rates between settings, and tested whether model-generated susceptibility measurements were associated with demographic and anthropometric factors. We identified greater variation in susceptibility to rotavirus infection and RVGE in Vellore than in Mexico City. In both cohorts, susceptibility to rotavirus infection and RVGE were associated with male sex, lower birth weight, lower maternal education, and having fewer siblings; within Vellore, susceptibility was also associated with lower socioeconomic status. Children who were more susceptible to rotavirus also experienced higher rates of diarrhea due to other causes. Simulations suggest that discrepant estimates of naturally-acquired immunity against RVGE can be attributed, in part, to between-setting differences in transmission intensity and susceptibility of children. We found that more children in Vellore than in Mexico City belong to a high-risk group for rotavirus infection and RVGE, and demonstrate that bias owing to differences in rotavirus transmission intensity and population susceptibility may hinder comparison of estimated immune protection against RVGE.

**Author summary:** Differences in susceptibility can help explain why some individuals, and not others, acquire infection and exhibit symptoms when exposed to infectious disease agents. However, it is difficult to distinguish between differences in susceptibility versus exposure in epidemiological studies. We developed a modeling approach to distinguish transmission intensity and susceptibility in data from cohort studies of rotavirus infection among children in Mexico City, Mexico, and Vellore, India, and evaluated how these factors may have contributed to differences in estimates of naturally-acquired immune protection between the studies. We found that more children were at “high risk” of acquiring rotavirus infection, and of experiencing gastroenteritis when infected, in Vellore versus Mexico City. The probability of belonging to this high-risk stratum was associated with recognized risk factors such as lower socioeconomic status, lower birth weight, and risk of diarrhea due to other causes. We also found the risk for rotavirus infections to cause symptoms declined with age, and was independent of acquired immunity. Together, these findings can account for estimates of lower protective efficacy of acquired immunity against rotavirus gastroenteritis in high-incidence settings, which mirrors estimates of reduced effectiveness of live oral rotavirus vaccines in low- and middle-income countries.

## Introduction

Rotavirus is the leading source of gastrointestinal disease burden in children globally, with nearly 10 million severe cases and 193,000 fatalities estimated to occur annually [1]. One decade after their rollout in high-income settings, live oral rotavirus vaccines are currently being introduced to national immunization programs of low- and middle-income countries (LMICs). However, randomized controlled trials and post-licensure studies have reported lower vaccine efficacy and effectiveness in LMICs compared to higher-income settings [2,3]. Understanding this performance gap is essential to maximizing the impact of rotavirus vaccines where they are needed most.

Recent observational studies have investigated how factors such as oral polio vaccine co-administration [4,5], exposure to breast milk antibodies [6], environmental enteropathy [7], and nutritional status [8,9] influence susceptibility of children to RVGE and performance of oral vaccines. Variation in susceptibility among individuals within and between studies—due to these or other unmeasured risk factors—constitutes a well known source of bias in estimates of vaccine efficacy and effectiveness [10–13]. Differential removal of highly-susceptible individuals to a (partially) “immune” state constitutes a form of frailty bias that may persist even in randomized studies [10,14]. While the possibility of such bias in rotavirus vaccine studies has been raised [15], assessing its contribution to variation in estimates of protection has been difficult. The distribution of risk factors across settings is not easily compared, and individual variation in susceptibility may be only partially attributable to known or measured risk factors.

Similarly-designed birth cohort studies undertaken in socioeconomically-distinct LMIC populations of Mexico City, Mexico and Vellore, India provide an opportunity to characterize heterogeneity in susceptibility to rotavirus infection and RVGE, and to assess its influence on estimates of immune protection [16,17]. While the two studies supplied similar estimates of naturally-acquired immune protection against re-infection, differences in estimates of protection against RVGE reflected discrepancies in estimated vaccine efficacy between Latin America and South Asia [18–20]. Whereas no children in Mexico City experienced moderate-to-severe RVGE after two or more previous infections, two previous infections were associated with only 57% protection against moderate-to-severe RVGE among children in Vellore [16,17]. Paired re-analysis of the studies has provided evidence that differences owe, in part, to the influence of a subset of “high-risk” individuals in the Vellore cohort—who experienced high rates of rotavirus infection and high risk for RVGE given infection—as well as age-dependent risk for RVGE given infection [21].

We revisited data from these studies aiming to better understand the distribution of susceptibility among individuals within the two cohorts, and to explore the implications for epidemiologic analyses. We developed a model to estimate susceptibility of children to rotavirus infection and RVGE, accounting for the natural history of rotavirus and differences between settings in transmission intensity. We used our findings to explore biases underlying conventional measures of protective immunity.

## Results

### Cohort monitoring

Incidence of rotavirus infection and RVGE among children enrolled in the two cohorts has been described previously [16,17,21,22]. Briefly, the studies enrolled 200 and 373 unvaccinated Mexican and Indian children who were followed from birth to up to 2 years and 3 years of age, respectively, yielding 3699 and 13,937 child-months of follow-up. In total, 315 rotavirus infections were detected in Mexico City and 1103 were detected in Vellore, with 89 (28% of 315 infections) and 282 (26% of 1103 infections) episodes of RVGE occurring in the two settings, respectively. Transmission intensity was higher in Vellore, such that first infections occurred in 56% and 81% of Indian children by ages 6 months and one year, compared to 34% and 67% of Mexican children, respectively (**Figs 1A, 1B**).

**Fig 1.**
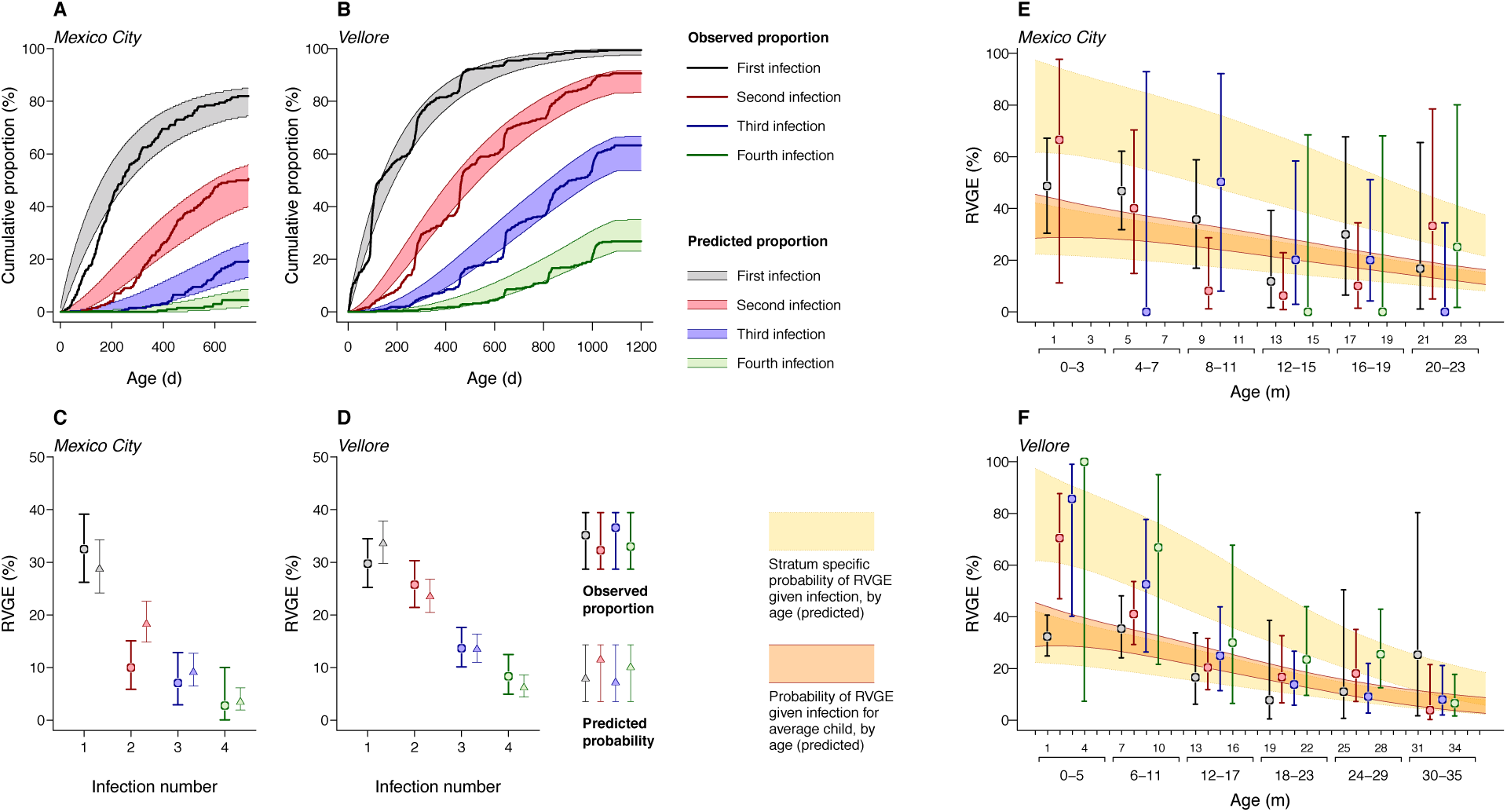
Natural history of rotavirus infection: observations and model predictions. (**A** and **B**) Cumulative incidence of first, second, third, and fourth rotavirus infections within the cohorts is plotted against model-based estimates; shaded areas define 95% prediction intervals as fitted by the model, accounting for variation in risk among children and for loss to follow-up. (**C** and **D**) Observed proportions (with 95% credible intervals) of infections involving RVGE, and model-predicted probabilities of RVGE given infection (with 95% prediction intervals), are plotted side by side. (**E** and **F**) Age-specific proportions of primary, secondary, third, and later infections involving RVGE (with 95% credible intervals) are plotted against model-estimated relationships between age and risk for infection to cause RVGE, within distinct risk strata (yellow; *R* and *R^C^*, see **Materials and Methods**) and for the average child in each cohort (orange; superimposed). Shaded areas delineate 95% prediction intervals around the age-specific probability. Age bins within which we calculate risk for RVGE are defined by the intervals between serological sampling dates within the two studies (4 months in Mexico City, 6 months in Vellore).

Simpson’s paradox was evident in the relationship among age, previous infection, and RVGE risk [21]. Whereas the proportion of infections causing RVGE declined with a higher number of previous infections (**Figs 1C, 1D**), this trend masked declining RVGE risk with older age for each of first, second, and later infections (**Figs 1E, 1F**). At matched ages, RVGE was more common during second and later infections than first infections in Vellore. In contrast, this trend was not apparent in Mexico City.

### Rotavirus natural history

We developed a model of rotavirus natural history guided by studies of transmission dynamics [23–25] and secondary analysis of the birth-cohort datasets [21]; our model also accounted for variation in susceptibility of children within the cohorts (see **Materials and Methods**). Fitting the model to the cohort datasets, we estimated 33% (95% CI: 23% to 41%), 50% (42% to 57%), and 64% (55% to 70%) reductions in the rates at which children re-acquired rotavirus after one, two, or three or more previous infections (**Table 1**), closely recapitulating estimated protection against re-infection in the original studies [16,17].

**Table 1:**
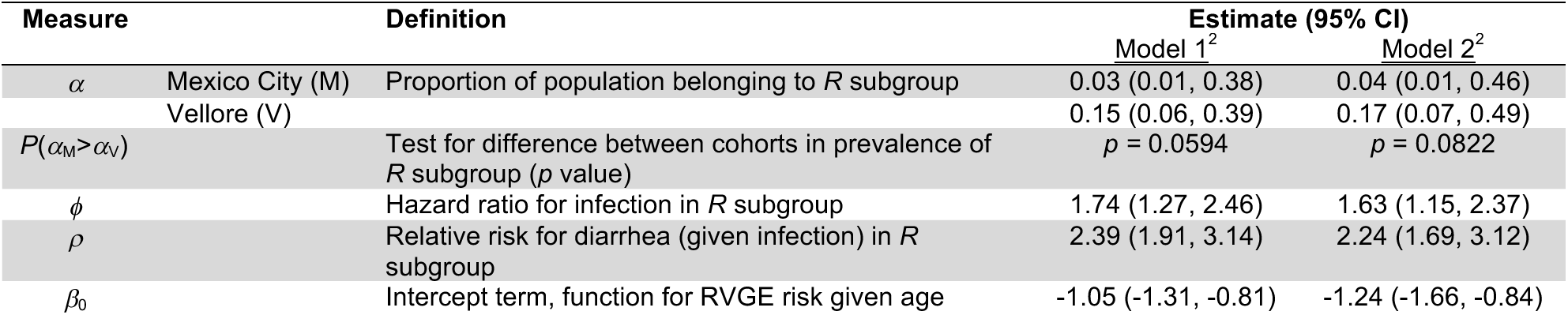
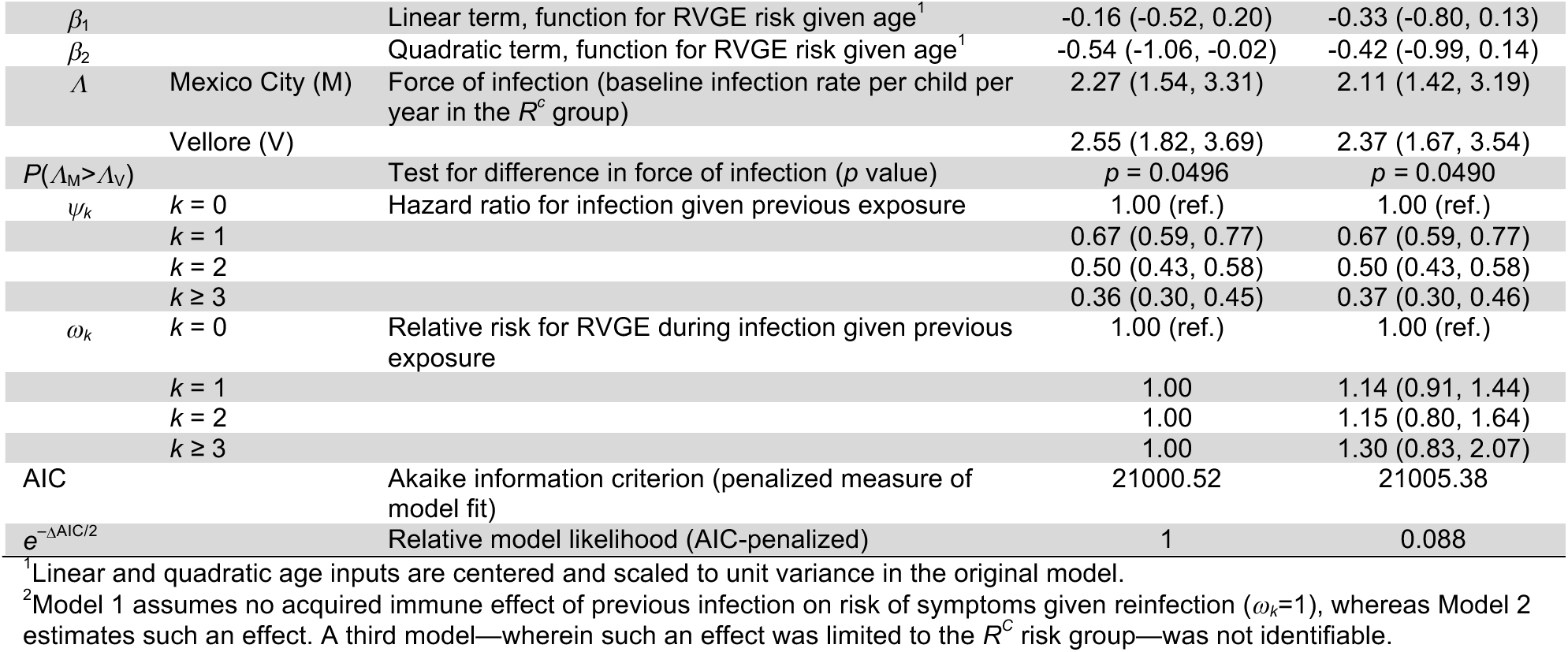
Parameter estimates and AIC values from models.

Our model also captured declining risk for infections to cause symptomatic RVGE at older ages (**Fig 1B**). The proportion of secondary and subsequent infections causing RVGE in the first year of life in Vellore closely matched expectations among children classified as a “high-risk” subset of the population (detailed below; **Fig 1F)**. We compared the fit of models with differing assumptions about acquired immune protection against RVGE given infection (**Table 1**). After accounting for declining risk of RVGE given infection at older ages, we did not identify improvements in model fit (based on values of the Akaike Information Criterion [26]) when allowing for acquired immune protection against symptoms during second or later infections.

Several salient differences between the two studies were reproduced in model-based predictions. Although the model predicted higher-than-observed rates of infection in Mexico City during the first six months of life, predictions accurately reflected between-setting differences in cumulative incidence by the end of the first year (**Figs 1A, 1B**). In addition, the model recapitulated the observation of significantly lower probabilities of RVGE during second, third, and fourth infections in Mexico City as compared to Vellore (**Figs 1C, 1D**), despite predicting RVGE in a higher-than-observed proportion of second infections in Mexico City.

### Variation in susceptibility among children

Our modeling framework partitioned the cohort populations across distinct risk groups (*R* and *R^C^*) with prevalences *α*_M_ and 1−*α*_M_, respectively, in Mexico City, and *α*_V_ and 1−*α*_V_ in Vellore. Because the size of the risk group and average relative risk are inversely related, the relative susceptibility and prevalence of these two risk groups were not simultaneously identifiable. We therefore estimated conditional between-group differences in susceptibility to infection (hazard ratio *ϕ*) and RVGE given infection (relative risk *ρ*) associated with particular values of *α*_M_ and *α*_V_. We reconstructed the full distribution of *ϕ* and *ρ* from the marginal distributions of {*ϕ,ρ*}|{*α*_M_,*α*_V_} (see **Materials and Methods**).

This modeling approach enabled us to compare the prevalence of children with particular susceptibility levels between cohorts (**Fig 2**). Defining a cut-off as a ≥50% higher-than-baseline rate of acquiring rotavirus infection, we estimated that 3% (1% to 23%) of children in Mexico City would belong to the high-risk stratum, compared to 13% (6% to 29%) of children in Vellore. A subgroup with over double the baseline hazard of infection would include 2% (1% to 8%) of children in Mexico City and 10% (5% to 18%) of children in Vellore, while if the relative rate of infection in the subgroup were more than three times the baseline hazard of infection, only 1% (0% to 2%) of children in Mexico City and 6% (5% to 9%) of children in Vellore would belong to the high-risk subgroup. Greater susceptibility to infection was associated with higher risk of experiencing RVGE given infection, regardless of the prevalence of the high-risk group (**Fig 2F**).

**Fig 2.**
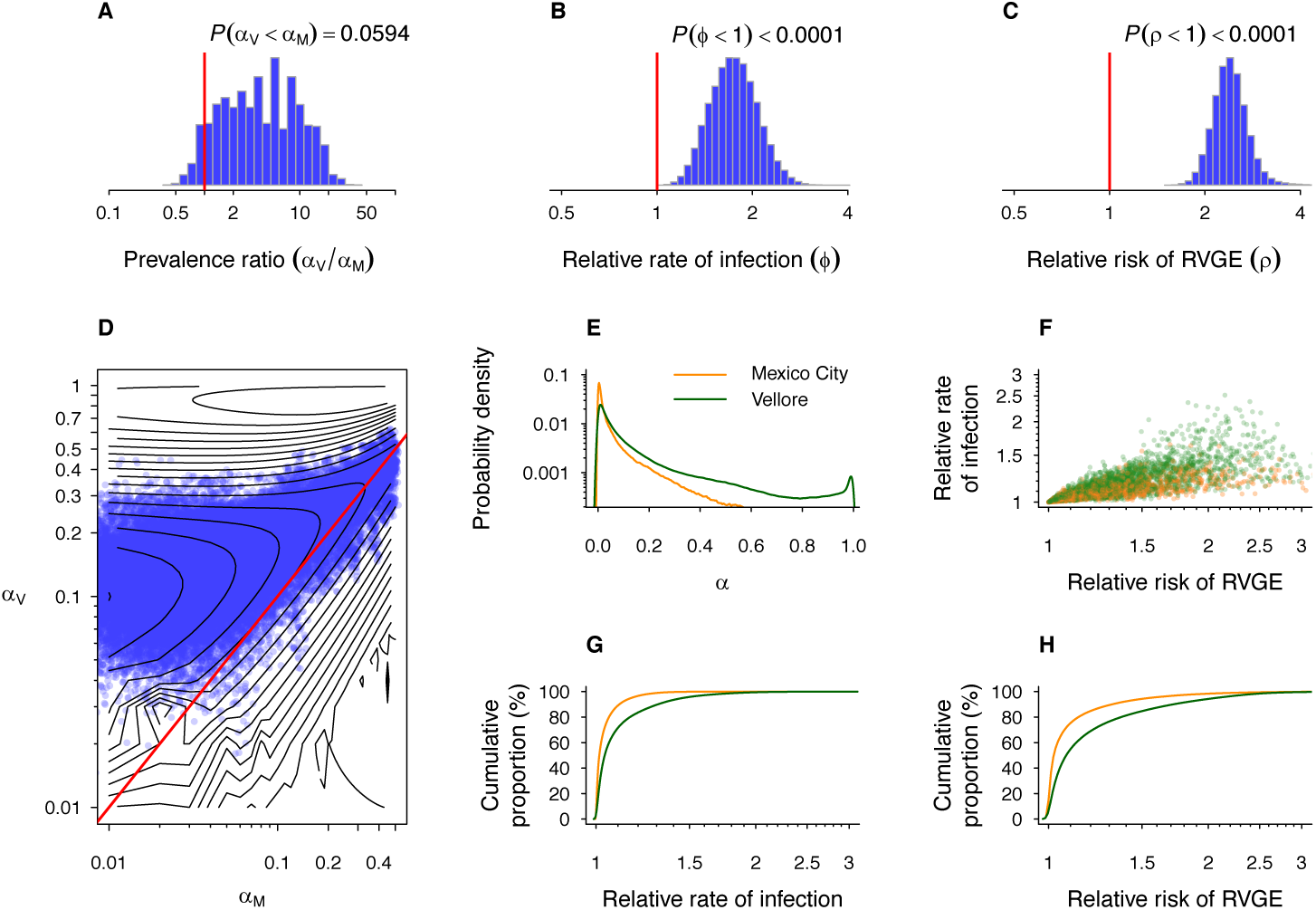
Heterogeneity in susceptibility to rotavirus infection and RVGE within and between cohorts. Histograms plot estimates of (**A** and **E**) the relative prevalence of a “higher risk” subgroup and (**B**) of subgroup-specific relative rates of infection and (**C**) relative risk of RVGE given infection. (**D**) Isoclines and sampled points (blue) illustrate the likelihood surface of the prevalence of subgroups within the Vellore and Mexico City cohorts. Integrating over all sampled possibilities for prevalence of the subgroups (**E**) and associated estimates of group-level differences in infection rate and RVGE risk supplies estimates of the relative susceptibility of individual children. At the level of individual children, estimates of susceptibility to infection are positively associated with susceptibility to RVGE when infected; points represent individual children sampled from the source populations according to parameter estimates (**F**). More variance in the distribution of acquisition rates (**G**) and RVGE risk (**H**) is evident in Vellore than Mexico City, signifying a greater prevalence of highly-susceptible children.

### Determinants of susceptibility among individual children

Our model provided a statistical basis for calculating the probability that each child belonged to the highrisk subgroup (see **Materials and Methods**). We used these estimated probabilities to examine the association between risk of belonging to the high-risk subgroup and host factors and exposures measured in the original studies (**Tables 2, 3**). We treated log-transformed probabilities as outcome variables in regression models, and adjusted for setting effects to account for differential prevalence of the risk factors in the two cohort populations.

**Table 2:** Characteristics of the two cohorts.

| Risk factor |  | Mexico City | Vellore |
| --- | --- | --- | --- |
| Birth weight | (kg) | 3.2 (median; IQR: 2.9, 3.4) | 3.0 (median; IQR: 2.7, 3.2) |
| Weight at 12 mo. | (kg) | 10.0 (median; IQR: 9.1, 10.6) | 7.9 (median; IQR: 7.3, 8.6) |
| Sex | Female | 94 (47.0%) | 186 (49.9%) |
|  | Male | 106 (53.0%) | 187 (50.1%) |
| Duration of breastfeeding<sup>1</sup> |  |  |  |
|  | <3 months | 59 (38.1%) | 15 (4.1%) |
|  | 3-5 months | 27 (17.4%) | 27 (7.3%) |
|  | 6-11 months | 31 (20.0%) | 56 (15.1%) |
|  | 12+ months | 38 (24.5%) | 272 (73.5%) |
| Number of siblings |  |  |  |
|  | 0 | 66 (33.8%) | 118 (31.6%) |
|  | 1 | 56 (28.7%) | 123 (33.0%) |
|  | 2 | 38 (19.5%) | 90 (24.1%) |
|  | 3+ | 35 (17.9%) | 42 (11.3%) |
| Socioeconomic status |  |  |  |
|  | Lowest | 24 (12.3%) | -- |
|  | Intermediate | 81 (41.5%) | -- |
|  | Highest | 90 (46.2%) | -- |
|  | Not bidi-working household | -- | 200 (53.6%) |
|  | Bidi-working household | -- | 173 (46.4%) |
| Maternal education<sup>2</sup> |  |  |  |
|  | 0-4 years | 57 (29.2%) | 106 (28.4%) |
|  | 5-7 years | 62 (31.8%) | 109 (29.2%) |
|  | 8-9 years | 46 (23.6%) | 90 (24.1%) |
|  | ≥10 years | 30 (15.4%) | 68 (18.2%) |
| Episodes of rotavirus-negative diarrhea<sup>3</sup> |  | 4 (median; IQR: 2, 7) | 4 (median; IQR: 2, 6) |
| Episodes of respiratory infection |  | -- | 20 (median; IQR: 14, 26) |

All percentages are calculated among children from whom data were available for the given variable. IQR: interquartile range.
<sup>1</sup>Calculated among children retained for over 1 year in the study.
<sup>2</sup>Breakpoints for the years of maternal education were defined according to the following standard for Vellore: 0-4, no formal education or primary school incomplete; 5-7, primary school complete; 8-9, middle school complete; 10 or more, high school complete.
<sup>3</sup>Duration of follow-up differs by setting (up to 2 years in Mexico City, versus up to 3 years in Vellore).

**Table 3:** Factors associated with belonging to “high-risk” group

| Risk factor |  | Relative Risk (95% CrI) |  |  |
| --- | --- | --- | --- | --- |
|  |  | Mexico City | Vellore | Pooled analysis |
| Birth weight | per log kg | **0.676 (0.418, 0.944)** | 0.841 (0.578, 1.129) | **0.486 (0.199, 0.977)** |
| Weight at 12 mo. | per log kg | 0.884 (0.740, 1.009) | 1.000 (0.845, 1.163) | 0.675 (0.234, 1.655) |
| Sex |  |  |  |  |
|  | Female | ref. | ref. | ref. |
|  | Male | 1.052 (0.788, 1.422) | **1.382 (1.079, 2.034)** | **1.264 (1.041, 1.666)** |
| Duration of breastfeeding<sup>1</sup> |  |  |  |  |
|  | <3 months | ref. | ref. | ref. |
|  | 3-5 months | 0.685 (0.407, 1.066) | 1.122 (0.434, 3.036) | 0.791 (0.495, 1.203) |
|  | 6-11 months | 0.913 (0.538, 1.438) | 1.158 (0.480, 2.860) | 0.912 (0.590, 1.367) |
|  | 12+ months | 0.698 (0.417, 1.053) | 1.110 (0.485, 2.510) | 0.830 (0.544, 1.166) |
| Number of siblings |  |  |  |  |
|  | 0 | ref. |  | ref. |
|  | 1 | 0.880 (0.593, 1.270) | **0.716 (0.455, 0.979)** | **0.761 (0.523, 0.971)** |
|  | 2 | **0.685 (0.419, 0.981)** | 0.792 (0.509, 1.155) | 0.753 (0.534, 1.001) |
|  | 3+ | **0.486 (0.282, 0.736)** | 0.742 (0.413, 1.118) | **0.606 (0.395, 0.833)** |
| Socioeconomic status |  |  |  |  |
|  | Lowest | ref. | -- | -- |
|  | Intermediate | 1.198 (0.777, 1.915) | -- | -- |
|  | Highest | 1.172 (0.772, 1.781) | -- | -- |
|  | Not bidi-working household | -- | ref. | -- |
|  | Bidi-working household | -- | **1.452 (1.095, 2.219)** | -- |
| Maternal education |  |  |  |  |
|  | 0-4 years (primary school incomplete) | ref. | ref. | ref. |
|  | 5-7 years (primary school complete) | 1.209 (0.864, 1.795) | 0.887 (0.621, 1.280) | 0.987 (0.758, 1.321) |
|  | 8-9 years (middle school complete) | 1.137 (0.771, 1.764) | 0.898 (0.612, 1.351) | 0.981 (0.733, 1.343) |
|  | ≥10 years (high school complete) | 1.128 (0.748, 1.798) | **0.619 (0.360, 0.902)** | **0.750 (0.502, 0.998)** |
| Incidence of rotavirus-negative diarrhea | per log episodes/year | **1.071 (1.022, 1.157)** | **1.376 (1.165, 1.715)** | **1.152 (1.072, 1.280)** |
| Incidence of respiratory infections | per log episodes/year | -- | **1.130 (1.104, 1.193)** | -- |

<sup>1</sup>Only children retained in studies for over 1 year are included in analyses.

Male children were 26% (4% to 67%) more likely to be among the high-risk subgroup than female children (**Table 3**). Birth weight was also a predictor being of in the high-risk group, with each log-kilogram decrease in birth weight conferring 2.05 (1.02 to 5.03)-fold higher probability of belonging to the high-risk subgroup. However, we did not detect a significant association between susceptibility and weight at 12 months. In each cohort, children without siblings were more likely to belong to the high-risk subgroup than children with siblings. In comparison to children whose mothers had completed <5 years of education, children whose mothers had completed ≥10 years of education were 25% (0% to 50%) less likely to belong to the high-risk subgroup.

We also identified several factors predicting within-cohort variation in susceptibility that were consistent with findings in primary analyses of the studies [16,17]. In Vellore, children whose household members were involved in producing bidis (indigenous cigarettes)—an indicator of lower household socioeconomic status—were 45% (10% to 122%) more likely than other children to belong to high-risk subgroup. In Mexico City, children with a shorter duration of breastfeeding were more likely to belong to the high-risk subgroup, although this association did not reach conventional thresholds of statistical significance in our analysis.

Children who experienced higher rates of diarrheal episodes caused by pathogens other than rotavirus were also more likely to belong to the high-risk subgroup within each cohort. In Vellore, we also found a positive association between the incidence of acute respiratory infections and the likelihood that a child belonged to the high-risk subgroup; this information was not available for the Mexico City cohort.

### Sources of variation in estimates of natural immune protection

We next conducted simulation studies (**Fig 3**) to assess how variation in transmission intensity and in the susceptibility of children could influence estimates of naturally-acquired immune protection against rotavirus infection, RVGE, and RVGE given infection [27]—as measured by the hazard ratios of infection and RVGE, and relative risk of RVGE given infection—following one, two, or three previous infections, compared to zero previous infections. We compared estimates from *in silico* cohorts with differing prevalence of “high risk” children exposed to varying forces of infection. We accounted for susceptibility differences between risk groups by sampling from the joint, unconditional distribution of *{ϕ,ρ}*, thereby isolating the effect of differences in risk-group prevalence.

**Fig 3.**
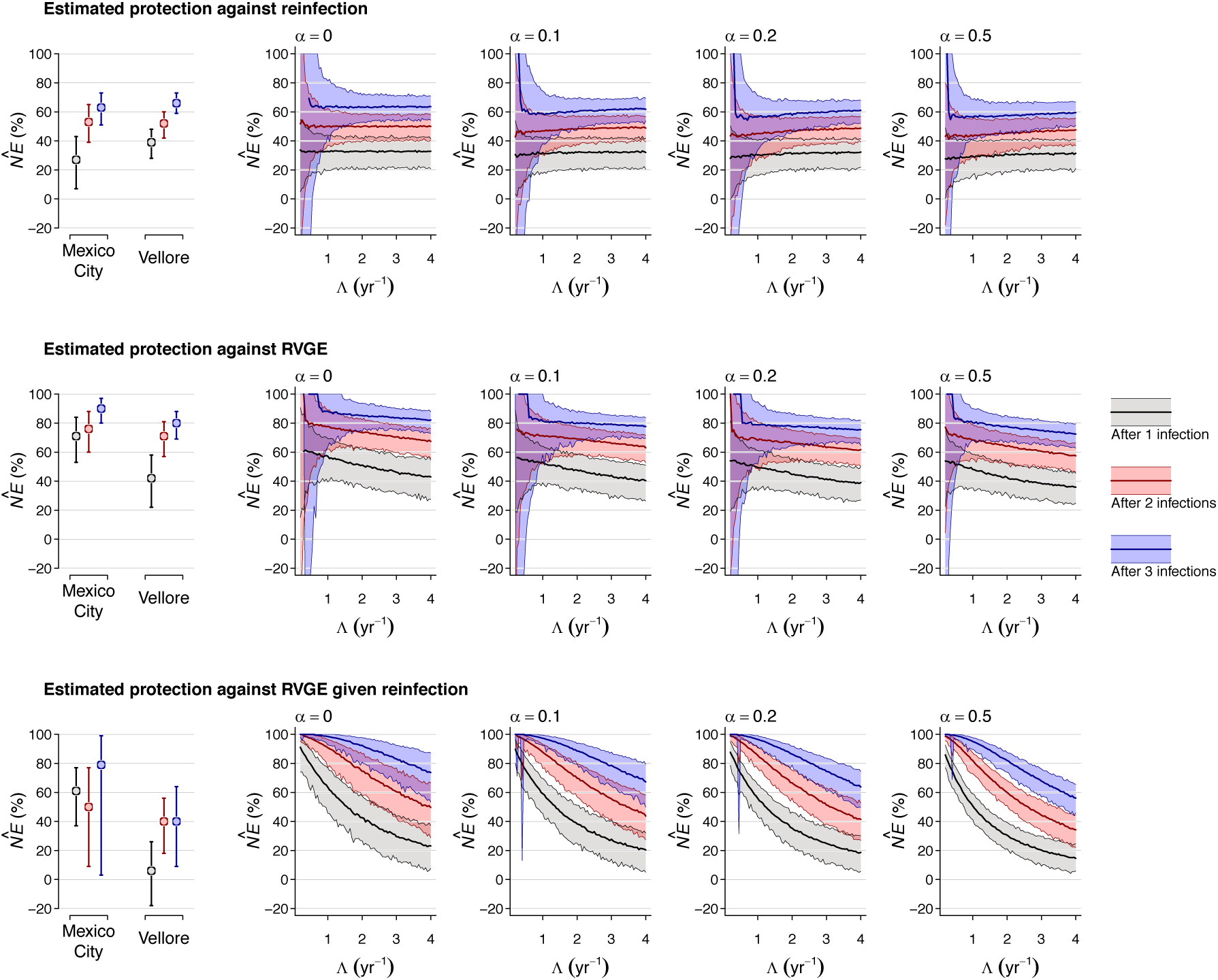
Sources of bias in estimates of naturally-acquired immune protection. We compared “naïve” estimates of naturally-acquired immunity against re-infection (top), against RVGE incidence (middle), and against RVGE given re-infection (bottom) in simulated cohorts using estimates of natural history parameters fitted from Model 1. We varied the force of infection and the proportion of children assumed to belong to a “high risk” group, sampling estimates of the relative rate of infection and relative risk of RVGE within this group from the unconditional distributions illustrated in **Figs 2B, 2C**. We plot estimates from the cohort datasets within the left panels of each row.

We did not identify a large degree of bias affecting estimates of protection against reinfection, which may help to explain why these estimates were nearly equal in the original studies [16,17]. However, we found that estimates of protection against RVGE—which were lower in primary analyses of the Vellore cohort—were expected to decline in settings with higher transmission intensity, reflecting acquisition of infection at younger, higher-risk ages. Moreover, this relationship was expected to be more pronounced in settings with a higher proportion of “high-risk” children (*α*).

## Discussion

Evidence of naturally-acquired immunity against rotavirus from birth-cohort studies provided an impetus toward the development of live oral rotavirus vaccines, which are now among the most effective strategies for the prevention of severe illness and deaths due to RVGE globally [28]. However, challenges have persisted in understanding and addressing the lower protective efficacy of rotavirus vaccines in high-burden LMIC settings, which mirrors protection derived from naturally-acquired immunity [29,30]. Our analysis suggests that discrepant estimates of protection may in part reflect epidemiological bias, attributable to differences between settings in transmission intensity and differential susceptibility of children to rotavirus infection and RVGE. Lower estimates of protection in settings with high rotavirus burden thus reflect factors other than weaker immunity among children in LMICs.

Accounting for aspects of the natural history of rotavirus enabled us to directly compare the susceptibility of children enrolled in birth cohort studies undertaken in socioeconomically-distinct settings. Although we estimated only modestly higher susceptibility for the average child in Vellore as compared to Mexico City, individual variation in susceptibility was considerably greater within the Vellore cohort. We estimated that a higher proportion of children in Vellore, as compared to Mexico City, showed elevated rates of rotavirus infection as well as excess risk for RVGE given infection. This finding can account for several unexpected features of the epidemiology of rotavirus in Vellore. First, increasing probability of RVGE in association with first, second, and later infections occurring at matched ages reflects high risk for RVGE given infection among individuals susceptible to frequent rotavirus infection. Indeed, our analysis identified that susceptibility to rotavirus infection was positively associated with susceptibility to RVGE given infection among individual children. Second, the proportion of children belonging to a high-risk group constituted a source of epidemiologic bias in simulation studies, and was expected to lead to estimates of weaker protection against RVGE in settings with higher transmission intensity such as Vellore.

While the influence of variation in susceptibility on rotavirus transmission dynamics and measures of intervention effectiveness have attracted methodological interest, optimal strategies to account for such variation in quantitative studies have remained elusive [12,13,15,31,32]. Directly comparing susceptibility between populations or settings is difficult because determinants of susceptibility are often unknown or unmeasured, and may be sub-optimally characterized by measurable epidemiological risk factors [14].

The contributions of susceptibility and transmission intensity to disease incidence rates are not easily disentangled, as incidence may reflect both of these factors. Our analysis employed a novel approach to characterize susceptibility of children in two cohorts based on a model that included known aspects of rotavirus natural history.

Our estimates of susceptibility are validated by their association with previously-reported risk factors for RVGE [33–36]: male children, children with lower birth weight, and children whose mothers had lower educational attainment were more likely to belong to a higher risk subset of the population in our analysis. In Vellore, children whose households were involved in bidi work—a marker for lower socioeconomic status—were also at higher risk [17], while in Mexico City, we observed a trend toward lower risk associated with longer breastfeeding, consistent with previous studies [37,38]. In addition, we observed higher incidence of diarrhea caused by pathogens other than rotavirus among children who were found to have greater susceptibility to rotavirus. This observation may signify the presence of environmental enteric dysfunction within the cohorts, or other sources of variation in immune status or pathogen exposure. In Vellore, children who we estimated were more susceptible to rotavirus also experienced higher incidence of respiratory infections, as reported previously [17].

These and other host factors associated with susceptibility to rotavirus infection and RVGE have also been reported to predict weaker immune responses to live enteric vaccines such as those against rotavirus. While our model does not address variation in the strength of immune responses, 58% of Indian children versus 90% of Mexican children seroconverted after Rotarix immunization in previous studies [29,39]. Nonetheless, near-equal naturally-acquired protection against re-infection was noted among children the birth cohort studies in Vellore and Mexico City. Our findings demonstrate that some degree of the reported variation in protection against RVGE can be attributed to epidemiological biases resulting from differential transmission intensity and differential susceptibility of children.

The finding that age diminishes risk for children to experience RVGE given rotavirus infection has been suggested in previous analyses of the cohort datasets [21]. Our simulation study demonstrates that such age-related symptom risk enhances protection against RVGE in low-transmission settings. Deferring infections to later ages significantly reduces the risk for children to experience symptoms upon reinfection. While the mechanisms underlying age-dependent diarrhea risk are not precisely known, the observation has been reported in mouse, rat, rabbit, and gnotobiotic piglet models of rotavirus infection [40–43]. Age-dependent TLR3 expression and host responses to rotavirus enterotoxins contribute to this observation in mice [44,45]. Other aspects of immune maturation, intestinal development, and the establishment of gut microbial communities may further drive associations between age and diarrhea risk in both humans and animals [46]. Furthermore, the greater dehydrating effect of diarrhea in younger children with smaller body volumes may contribute to severity—and thus the reporting and diagnosis—of RVGE in early-life infections [47].

There are several limitations to our analysis. Whereas we assume exponentially-distributed infection times (consistent with a constant hazard of infection), this provides an imperfect fit to the timing of early-life infections, particularly in the Mexico City cohort. The departure between predictions and observations may reflect the protective effect of maternal antibodies, as reported previously [16,48], or the influence of age-specific social mixing patterns on transmission [49]. Thus, our model tended to overestimate the probability of RVGE associated with second rotavirus infections in Mexico City, although this discrepancy was not sustained for third and fourth infections. Our analyses also do not distinguish between homotypic and heterotypic protection because we lack genotype data for serologically-detected infections, which constitute the majority of infections in both cohorts. Although moderate-to-severe RVGE episodes are the primary endpoint of most studies evaluating vaccine efficacy and effectiveness, our analysis addressed RVGE episodes of any severity. Only eight RVGE episodes with Vesikari score ≥11 were observed in Mexico City, limiting our statistical power for analyses of moderate-to-severe RVGE. Nonetheless, previous analyses of the studies identified similar risk factors for mild and moderate-to-severe RVGE [21], suggesting our findings can also inform the interpretation of studies with moderate-to-severe RVGE endpoints.

Birth-cohort studies have been instrumental to our understanding of the natural history of rotavirus. Uncertainties surrounding differences in the epidemiology of rotavirus in socioeconomically-distinct populations underscore the need for a theoretical basis for comparing outcomes of individual studies. Our approach permitted assessment of how age, acquired immunity, and predictors of individual susceptibility independently contributed to infection and disease risk in distinct birth cohorts, resolving discrepancies in estimated protection that arose in primary analyses of the datasets. The modeling framework we introduce here may thus have applicability to studies of other partially-immunizing pathogens.

## Materials and methods

### Birth cohort data

The two birth cohort studies followed similar protocols that have been described previously [16,17]. Children were enrolled at birth and followed to 24 and 36 months of age in Mexico City and Vellore, respectively. The studies aimed to detect all rotavirus infections, both symptomatic and asymptomatic. Rotavirus infections were detected by three approaches: (1) sera were drawn every 4 and 6 months in Mexico City and Vellore, respectively, and tested for IgA or IgG titer increases; (2) asymptomatic stool samples were collected weekly in Mexico City and every two weeks in Vellore and tested for rotavirus; and (3) diarrheal stools were collected by field workers every time mothers alerted the study teams of any change in a child’s stool pattern (**S1 Table**). Virus detection was performed by ELISA in Mexico City and by ELISA or real-time PCR in Vellore. In Mexico City, 200 children were recruited and retained for 77% of the scheduled follow-up period, while our analysis of the Vellore dataset included the 373 children (83% of 452 enrolled) who completed follow-up. Data were available for 96% (1037/1080) and 99% (2565/2598) of scheduled serum tests; 97% (15,503/16,029) and 93% (26,902/28,906) of scheduled asymptomatic stool tests; and 85% (963/1133) and 99% (1829/1856) of reported diarrheal episodes in Mexico City and Vellore, respectively.

### Modeling rotavirus natural history

**Summary**. We developed a probabilistic model describing rotavirus natural history that allowed us to test how susceptibility to rotavirus infection and RVGE vary innately among individuals and according to age and previous infection. Seeking to account for differences in rotavirus epidemiology in the two settings under a unified model, we assumed parameters reflecting the strength of immune protection and the effect of age on the probability of symptoms given infection were equal in both settings, whereas we permitted the force of infection and the proportion of children belonging to distinct risk strata to vary between settings. We compared the improvement in fit afforded by modifications reflecting distinct hypotheses about the acquisition of protection against symptoms given infection—a source of uncertainty in rotavirus epidemiology [21]. Individual-level susceptibility of children to infection and RVGE was measured from the probability for children to belong to the high-risk stratum, which we also tested for associations with epidemiological risk factors. Last, we simulated from the fitted model to explore how differences in transmission intensity and the prevalence of high-risk individuals influence estimates of protection.

**Model**. Our model of rotavirus natural history accounted for the rate at which individuals acquired and re-acquired rotavirus infection, as well as their probability of experiencing RVGE when infected.

Observations from previous analyses of the cohort studies suggested individuals differ in their susceptibility: in Vellore, children who experienced diarrhea on second and later rotavirus infections were more likely to have experienced diarrhea on earlier-life infections [21]. We therefore accounted for potential variation in risk among children by defining proportions *α*_V_ and α_M_ of children in Vellore and Mexico City, respectively, belonging to a distinct risk group *R*, and tested for group-wise differences in susceptibility to rotavirus infection and RVGE in the *R* stratum relative to the remaining population (*R^c^*, with prevalence 1−*α*_s_ in setting *s*). To avoid generating symmetric estimates of lower and higher risk groups, we defined *α*_M_≤0.5 and 0≤*α*_V_≤1; while this presumed the *R* population would constitute a higher-risk group, this designation formally depended upon estimates of the relative susceptibility of children (**Table 1**).

Whereas primary analyses of the cohort datasets identified decreasing RVGE incidence rates following previous rotavirus infection, re-analyses suggested this declining risk resulted from naturally-acquired protection against rotavirus infection and older age at time of reinfection [21]. We used the Akaike Information Criterion (AIC) [26] to compare the fit of alternative, nested models premised on differing assumptions about the influence of age and naturally-acquired immunity on risk of experiencing diarrhea symptoms during rotavirus infection. Model 1 assumed the risk of RVGE given infection did not vary based on the number of previous infections. Model 2 considered both age and previous infection as determinants of risk for RVGE given infection. We also attempted to fit a third model assuming that only children in the *R^C^* (effectively, lower-risk) group acquired protection against symptoms given infection, but found that parameters were not identifiable; thus, only Models 1 and 2 were considered.

We modeled infection at the rate 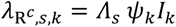 among children in the majority (*R^c^*) risk group, where *Λ*_s_ was the force of infection in Mexico City or Vellore, *I_k_* was an indicator that an individual had experienced *k* previous rotavirus infections, and *ψ_k_* was the hazard ratio for reinfection resulting from naturally-acquired immune protection following *k* previous infections. Among children in the minority (*R*) risk group, we modeled 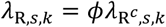, where the hazard ratio *ϕ* conveyed the effect of differential susceptibility to infection.

The probability of RVGE given rotavirus infection, *π*, was modeled as a function of age and risk group under Model 1. For children of age *x* in the majority risk group,

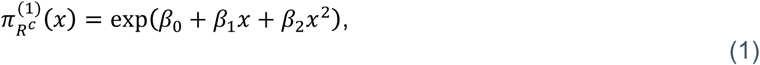
where the parameters *β*_0–2_ related RVGE risk to the child’s age. Linear and higher-order polynomial functions relating risk to age were explored, but found not to improve model fit via AIC. For the remaining children,

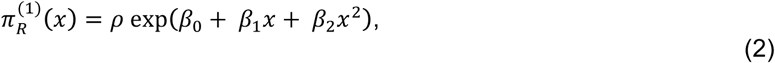
where the relative risk *ρ* accounted for innate differences in susceptibility to RVGE given infection.

Under Models 2 and 3, we further accounted for variation in symptom risk according to the number of previous infections a child in the *R^c^* stratum has experienced, such that for Model 2,

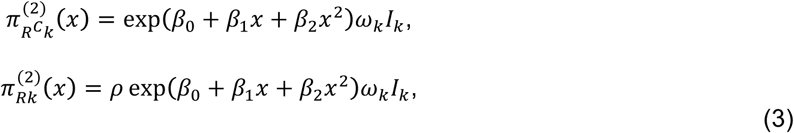
where *ω_k_* indicated the fold change in risk of RVGE given infection after having experienced *k* previous infections. For Model 3,

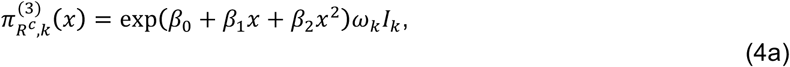
whereas

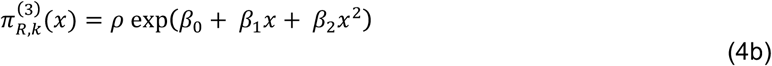
for all *k*.

**Likelihood**. We estimated model parameters in a likelihood-based framework. Observations included all infections or instances of censoring (indexed by *j*) for each child (indexed by *i*). For the *j*th observation of the *i*th child, we denoted time since the last observation or birth as *Δt_ij_*, indicating whether the child was currently infected with rotavirus with the variable *Z_ij_* (=1 if infected and 0 if censoring occurred) and experiencing RVGE with the variable *D_ij_* (=1 if RVGE occurred upon infection and 0 otherwise). The likelihood contribution of observation *j* from child *I*, conditioned on the child belonging to either the *R* or *R^c^* subgroup (*) and residing in setting *s*, was

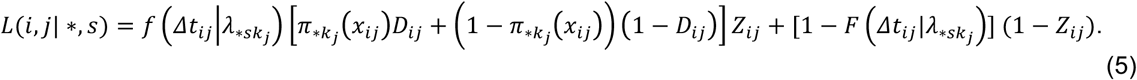

The term 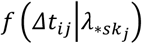 represents the probability that the time to infection for individual *I* was *Δt_ij_* while 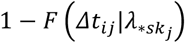 was the probability for child *i* to have escaped infection from the time of the previous observation to the last follow-up visit, if censoring occurred at the *j*th observation. We assumed an exponentially-distributed time to infection, consistent with a constant force of infection.

We obtained the likelihood contribution of each child, *H_s_*(*i*), via the total probability

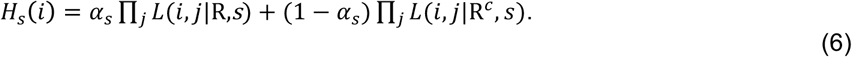

Defining **Y*_j_***, as the set of observations from each child *i*, the probability for any particular child to be in the minority risk (R) group was

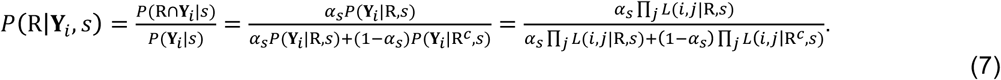

Our statistical framework assumed that children were sampled at random from the population and that observations **Y***_i_* are randomly-assigned given the setting, age, risk group, and infection history of an individual child *i*, such that the sample proportion of children in the *R* group was expected to converge to the population proportion.

We thus defined the overall model likelihood in two components. The first, 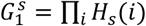, conveyed the direct contributions of observations **Y***_i_* as described above. The second component, 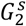, reflected concordance between the population proportion, *α*_s_, and the sample proportion, 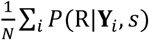 of children in the *R* risk group. We calculated 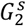 by evaluating the probability density of *α_s_* under the assumption of random assignment:

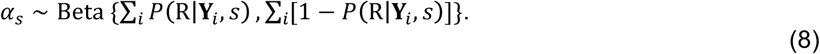

Altogether, the model likelihood was the product of these terms across the two settings, 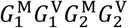.

**Estimation**. Estimated between-group differences in susceptibility to infection and disease (*ϕ* and *ρ*, respectively) were dependent upon the size of the underlying proportions *α*_M_ and *α*_V_. We therefore conducted inference via the conditional distributions of the model parameters *θ =* {*ϕ*, *ρ, ψ_1–3_, ω_1–3_, β_0–2_*, *Λ_m_, Λ_v_*} over fixed values of {*α*_M_, *α*_V_}, defined over [0, 0.01,…, 0.5] and [0, 0.01,…, 1], respectively.

For each set {*α*_M_, *α*_V_}, we determined the conditional maximum likelihood parameter estimates *θ*^*^|*α*_M_*, α*_V_ via the Nelder-Mead algorithm:

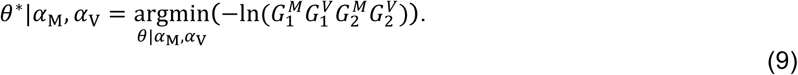

We accounted for parameter uncertainty at each set {*α*_M_, *α*_V_} by sampling from the multivariate normal distributions surrounding conditional maximum-likelihood estimates. We pooled these draws over all {*α*_M_, *α*_V_}, weighting by their respective likelihoods, to recover the unconditional parameter distributions.

**Determinants of individual risk**. To better understand variation in susceptibility among children within each cohort, we evaluated associations between individual-level factors (**Table 2**) and the probability for each child to belong to the high-risk (*R*) subgroup. For each of 10,000 draws of *θ*, we measured the probability of belonging to the high-risk group, equal to P(R|**Y***_i_*, *s*) for *ρ*>1 or 1 − *P*(R|**Y***_i_*, *s*) for *p*<1, for each child (in all samples, we identified *ϕ*>1 for *ρ*>1 and *ϕ*<1 for *ρ*<1). We used least-squares regression to test for associations between covariates and children’s log-transformed probability of being in the high-risk group, using estimated regression parameters to measure relative risks (**Table 3**). Models included a setting term to account for differential prevalence of high-risk children. We pooled relative risk estimates across our draws of *θ* to recover their distribution.

### Simulation study

To examine potential bias in conventional estimates of naturally-acquired immune protection, we used our model of the natural history of rotavirus infection to simulate individual histories of infection and RVGE over the first three years of life, sampling from estimated parameters describing the effects of age (*β*_0_, *β*_1_, *β*_2_) and previous infection (*ψ*_1_, *ψ*_2_, *ψ*_3_) on susceptibility to RVGE and infection, respectively. We conducted simulations under an external force of infection (*Λ*) ranging from 0.2 to 4 infections per year, assigning 0% to 50% (*α*) of children to the high-risk subgroup *R*; values of *ϕ* and *ρ* were drawn independently of *α* so that we could determine the effect of differences in the proportion of high-risk children on estimates of protection. We sampled exponentially-distributed infection times (calculated from time of birth or previous infection), and defined the occurrence of RVGE for each individual infection as a Bernoulli random variable using the model-predicted probability of RVGE given infection.

For each cohort simulation, we measured the hazard ratio for reinfection and RVGE from the incidence rate *(IR_k_)* of infection and RVGE after one, two, or three previous infections, relative to the *IR*_0_ from birth. We also measured the relative risk of RVGE given reinfection among children who had experienced one, two, or three previous infections, relative to those with no previous infections, calculated from the proportion (*p_k_*) of infections with RVGE. We defined estimates of natural immune efficacy 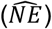 as

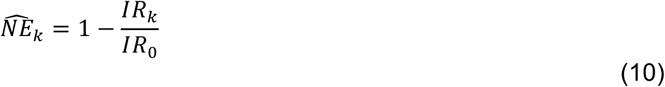
for protection against infection and RVGE among children who had experienced *k* previous infections, and

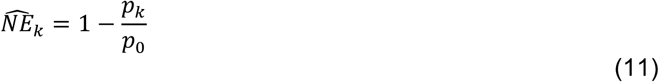
for RVGE progression among children who had experienced *k* previous infections.

---

**Table S1.** Study design, enrollment, and follow-up.

|  | **Mexico City cohort** | **Vellore cohort** |
| --- | --- | --- |
| Duration of follow-up | 24 months | 36 months |
| Frequency of asymptomatic stool testing | Weekly | Every 2 weeks |
| Frequency of serological testing | Every 4 months | At least every 6 months |
| Definition of rotavirus shedding | ELISA positive | 2x ELISA positive or RT-PCR positive |
| Definition of seroconversion | 4-fold rise in IgG or IgA | 4-fold rise in IgG or 3-fold rise in IgA |
| Study population | 200 children | 373 children<sup>1</sup> |
| Child-months of observation | 3699/4800 (77%) | 13341/13428 (99%) |
| Asymptomatic stool samples tested | 15503 | 26902 |
| Of all scheduled tests | 15503/20800 (75%) | 26902/29094 (92%) |
| Of all scheduled tests while child was retained in follow-up | 15503/16029 (97%) | 26902/28906 (93%) |
| Diarrheal episodes tested, of all reported diarrheal episodes | 963/1133<sup>2</sup> (85%) | 1829/1856 (99%) |
| Serum samples tested, of all scheduled tests | 1037/1080 (96%) | 2565/2598 (99%) |
| Infections detected | 316 | 1103 |
| From diarrheal episodes | 89/316 (28%) | 282/1103 (26%) |
| From asymptomatic shedding | 88/316 (28%) | 237/1103 (21%) |
| From seroconversion only | 139/316 (44%) | 584/1103 (53%) |
| Mean age (d), asymptomatic infections detected by seroconversion only | 445 (SD=167)<sup>3</sup> | 626 (SD=308)<sup>4</sup> |
| Mean age (d), asymptomatic infections detected by shedding | 339 (SD=187)<sup>3</sup> | 542 (SD=342)<sup>4</sup> |
| Rotavirus-negative diarrhea samples | 874/963 (91%) | 1547/1829 (85%) |

<sup>1</sup>Sample restricted to children who completed 3 years of follow-up (83% of initial cohort of 452 children)
<sup>2</sup>The total number was not provided in the original study, but was calculated from the information that the 963 tested episodes represented 85% of the total reported episodes.
<sup>2</sup>In the Mexico City cohort, the mean age of asymptomatic infections detected by shedding is lower than the mean age of asymptomatic infections detected by seroconversion alone ( $p<0.0001$ ).
<sup>3</sup>In the Vellore cohort, the mean age of asymptomatic infections detected by shedding is lower than the mean age of asymptomatic infections detected by seroconversion alone ( $p<0.001$ ).

## REFERENCES

1. Fischer Walker CL, Rudan I, Liu L, Nair H, Theodoratou E, Bhutta Z a., et al. Global burden of childhood pneumonia and diarrhoea. Lancet. 2013;381: 1405-1416.

2. Parashar UD, Johnson H, Steele AD, Tate JE. Health impact of rotavirus vaccination in developing countries: Progress and way forward. Clin Infect Dis. 2016;62: S91-S95.

3. Jiang V, Jiang B, Tate J, Parashar UD, Patel MM. Performance of rotavirus vaccines in developed and developing countries. Hum Vaccin. 2010;6: 532-42.

4. Kirkpatrick BD, Colgate ER, Mychaleckyj JC, Haque R, Dickson DM, Carmolli MP, et al. The “Performance of Rotavirus and Oral Polio Vaccines in Developing Countries” (PROVIDE) study: Description of methods of an interventional study designed to explore complex biologic problems. Am J Trop Med Hyg. 2015;92: 744-751.

5. Patel M, Steele AD, Parashar UD. Influence of oral polio vaccines on performance of the monovalent and pentavalent rotavirus vaccines. Vaccine. 2012;30: S30-S35.

6. Ali A, Kazi AM, Cortese MM, Fleming JA, Moon SS, Parashar UD, et al. Impact of withholding breastfeeding at the time of vaccination on the immunogenicity of oral rotavirus vaccine - A randomized trial. PLoS One. 2015;10:e0127622.

7. Mohan VR, Ramanujam K, Babji S, McGrath M, Shrestha S, Shrestha J, et al. Rotavirus infection and disease in a multi-site birth cohort: Results from the MAL-ED study. J Infect Dis. 2017;216: 305-316.

8. Perez-Schael I, Salinas B, Tomat M, Linhares AC, Guerrero ML, Ruiz-Palacios GM, et al. Efficacy of the human rotavirus vaccine RIX4414 in malnourished children. J Infect Dis. 2007;196: 537-540.

9. Levine MM, Kotloff KL, Nataro JP, Muhsen K. The Global Enteric Multicenter Study (GEMS): Impetus, rationale, and genesis. Clin Infect Dis. 2012;55: S215-S224.

10. Halloran ME, Longini IM, Struchiner CJ. Estimability and interpretation of vaccine efficacy using frailty mixing models. Am J Epidemiol. 1996;144: 83-97.

11. Smith PG, Rodrigues LC, Fine PEM. Assessment of the protective efficacy of vaccines against common diseases using case-control and cohort studies. Int J Epidemiol. 1984;13: 87-93.

12. Greenwood M, Yule GU. An inquiry into the nature of frequency distributions representative of multiple happenings with particular reference to the occurrence of multiple attacks of disease or of repeated accidents. J R Stat Soc. 1920;83: 255-279.

13. Hernán MA. The hazards of hazard ratios. Epidemiology. 2010;21: 13-15.

14. Morozova O, Cohen T, Crawford FW. Risk ratios for contagious outcomes. arXiv. 2017; Available: http://arxiv.org/abs/1707.05884.

15. Gomes MGM, Gordon SB, Lalloo DG. Clinical trials: The mathematics of falling vaccine efficacy with rising disease incidence. Vaccine. 2016;34: 3007-3009.

16. Velázquez FR, Matson DO, Calva JJ, Guerrero L, Morrow AL, Carter-Campbell S, et al. Rotavirus infections in infants as protection against subsequent infections. N Engl J Med. 1996;335: 1022-1028.

17. Gladstone BP, Ramani S, Mukhopadhya I, Muliyil J, Sarkar R, Rehman AM, et al. Protective effect of natural rotavirus infection in an Indian birth cohort. N Engl J Med. 2011;365: 337-346.

18. Ruiz-Palacios GM, Perez-Schael I, Velazquez FR, Abate H, Breuer T, Clemens SC, et al. Safety and efficacy of an attenuated vaccine against severe rotavirus gastroenteritis. N Engl J Med. 2006;354: 11-22.

19. Zaman K, Anh DD, Victor JC, Shin S, Yunus M, Dallas MJ, et al. Efficacy of pentavalent rotavirus vaccine against severe rotavirus gastroenteritis in infants in developing countries in Asia: A randomised, double-blind, placebo-controlled trial. Lancet. 2010;376: 615-623.

20. Bhandari N, Rongsen-Chandola T, Bavdekar A, John J, Antony K, Taneja S, et al. Efficacy of a monovalent human-bovine (116E) rotavirus vaccine in Indian infants: A randomised, double-blind, placebo-controlled trial. Lancet. 2014;383: 2136-2143.

21. Lewnard JA, Lopman BA, Parashar UD, Bar-Zeev N, Samuel P, Guerrero ML, et al. Naturally-acquired immunity against rotavirus infection and gastroenteritis in children: paired re-analyses of birth-cohort studies. J Infect Dis. 2017;216: 317-326.

22. Raul Velazquez F, Calva JJ, Lourdes Guerrero M, Mass D, Glass RI, Pickering LK, et al. Cohort study of rotavirus serotype patterns in symptomatic and asymptomatic infections in Mexican children. Pediatr Infect Dis J. 1993;12: 54-61.

23. Pitzer VE, Viboud C, Simonsen L, Steiner C, Panozzo CA, Alonso WJ, et al. Demographic variability, vaccination, and the spatiotemporal dynamics of rotavirus epidemics. Science. 2009;325: 290-294.

24. Pitzer VE, Atkins KE, de Blasio BF, van Effelterre T, Atchison CJ, Harris JP, et al. Direct and indirect effects of rotavirus vaccination: Comparing predictions from transmission dynamic models. PLoS One. 2012;7: e42320.

25. Atkins KE, Shim E, Carroll S, Quilici S, Galvani AP. The cost-effectiveness of pentavalent rotavirus vaccination in England and Wales. Vaccine. 2012;30: 6766-6776.

26. Akaike H. A new look at the statistical model identification. IEEE Trans Automat Contr. 1974;19: 716-723.

27. Halloran ME, Struchiner CJ, Longini IM. Study designs for evaluating different efficacy and effectiveness aspects of vaccines. Am J Epidemiol. 1997;146: 789-803.

28. Patel MM, Steele D, Gentsch JR, Wecker J, Glass RI, Parashar UD. Real-world impact of rotavirus vaccination. Pediatr Infect Dis J. 2011; 30: S1-S5.

29. Patel M, Shane AL, Parashar UD, Jiang B, Gentsch JR, Glass RI. Oral rotavirus vaccines: how well will they work where they are needed most? J Infect Dis. 2009;200: S39-S48.

30. Lopman BA, Pitzer VE, Sarkar R, Gladstone B, Patel M, Glasser J, et al. Understanding reduced rotavirus vaccine efficacy in low socio-economic settings. PLoS One. 2012;7: e41720.

31. Ball F. Deterministic and stochastic epidemics with several kinds of susceptibles. Adv Appl Probab. 1985;17: 1-22.

32. O’Hagan JJ, Hernán MA, Walensky RP, Lipsitch M. Apparent declining efficacy in randomized trials: examples of the Thai RV144 HIV vaccine and South African CAPRISA 004 microbicide trials. AIDS. 2012;26: 123-126.

33. Dennehy PH, Cortese MM, Bégué RE, Jaeger JL, Roberts NE, Zhang R, et al. A case-control study to determine risk factors for hospitalization for rotavirus gastroenteritis in US children. Pediatr Infect Dis J. 2006;25: 1123-1131.

34. Newman RD, Grupp-Phelan J, Shay DK, Davis RL. Perinatal risk factors for infant hospitalization with viral gastroenteritis. Pediatrics. 1999;103: 1-6.

35. Victoria CG, Huttly SRA, Barros FC, Lombardi C, Vaughan JP. Maternal education in relation to early and late child health outcomes: Findings from a Brazilian cohort study. Soc Sci Med. 1992;34: 899-905.

36. Naficy AB, Abu-Elyazeed R, Holmes JL, Rao MR, Savarino SJ, Kim Y, et al. Epidemiology of rotavirus diarrhea in Egyptian children and implications for disease control. Am J Epidemiol. 1999;150: 770-777.

37. Plenge-Bönig A, Soto-Ramírez N, Karmaus W, Petersen G, Davis S, Forster J. Breastfeeding protects against acute gastroenteritis due to rotavirus in infants. Eur J Pediatr. 2010;169: 1471-1476.

38. Newburg DS, Peterson JA, Ruiz-Palacios GM, Matson DO, Morrow AL, Shults J, et al. Role of human-milk lactadherin in protection against symptomatic rotavirus infection. Lancet. 1998;351: 1160-1164.

39. Patel M, Glass RI, Jiang B, Santosham M, Lopman B, Parashar U. A systematic review of anti-rotavirus serum IgA antibody titer as a potential correlate of rotavirus vaccine efficacy. J Infect Dis. 2013;208: 284-294.

40. Ciarlet M, Conner ME, Finegold MJ, Estes MK. Group A rotavirus infection and age-dependent diarrheal disease in rats: a new animal model to study the pathophysiology of rotavirus infection. J Virol. 2002;76: 41-57.

41. Ciarlet M, Gilger M a, Barone C, McArthur M, Estes MK, Conner ME. Rotavirus disease, but not infection and development of intestinal histopathological lesions, is age restricted in rabbits. Virology. 1998;251: 343-60.

42. Ramig RF. The effects of host age, virus dose, and virus strain on heterologous rotavirus infection of suckling mice. Microb Pathog. 1988;4: 189-202.

43. Saif LJ, Ward LA, Yuan L, Rosen BI, To TL. The gnotobiotic piglet as a model for studies of disease pathogenesis and immunity to human rotaviruses. Arch Virol. 1996;12: 153-161.

44. Pott J, Stockinger S, Torow N, Smoczek A, Lindner C, McInerney G, et al. Age-dependent TLR3 expression of the intestinal epithelium contributes to rotavirus susceptibility. PLoS Pathog. 2012;8: e1002670.

45. Ball JM, Tian P, Zeng CQ, Morris AP, Estes MK. Age-dependent diarrhea induced by a rotaviral nonstructural glycoprotein. Science. 1996;272: 101-4.

46. Fulde M, Hornef MW. Maturation of the enteric mucosal innate immune system during the postnatal period. Immunological Reviews. 2014;206: 21-34.

47. Cohen MB. Etiology and mechanisms of acute infectious diarrhea in infants in the United States. J Pediatr. 1991; 118: S34-S39.

48. Velázquez FR, Matson DO, Guerrero ML, Shults J, Calva JJ, Morrow AL, et al. Serum antibody as a marker of protection against natural rotavirus infection and disease. J Infect Dis. 2000;182: 1602-1609.

49. Mossong J, Hens N, Jit M, Beutels P, Auranen K, Mikolajczyk R, et al. Social contacts and mixing patterns relevant to the spread of infectious diseases. PLoS Med. 2008;5: e74.

